# Proteome Integral Stability Alteration assay dramatically increases throughput and sensitivity in profiling factor-induced proteome changes

**DOI:** 10.1101/496398

**Authors:** Massimiliano Gaetani, Pierre Sabatier, Amir Ata Saei, Christian Beusch, Zhe Yang, Susanna Lundström, Roman A. Zubarev

**Affiliations:** Division of Physiological Chemistry I, Department of Medical Biochemistry and Biophysics, Karolinska Institutet, SE-17 177 Stockholm, Sweden; SciLIfeLab, SE-17 177 Stockholm, Sweden; Department of Pharmacological & Technological Chemistry, I.M. Sechenov First Moscow State Medical University, Moscow, 119146, Russia

## Abstract

Various factors, including drugs as well as non-molecular influences, induce alterations in the stability of proteins in cell lysates, living cells and organisms. These alterations can be probed by applying a stability-modifying agent, such as elevated temperature, to a varying degree. As a second dimension of variation, drug concentration or factor intensity can be used. However, the corresponding analysis scheme has a low throughput and high cost. Additionally, since traditional data analysis employs curve fitting, proteins with unusual behavior are frequently ignored. The novel Proteome Integral Stability Alteration (PISA) assay avoids these issues altogether, increasing the analysis throughput by one to two orders of magnitude for unlimited number of parameter variation points. The consumption of the compound and biological material decreases by the same factor. We envision widespread use of the PISA approach in chemical biology and drug development.

## Introduction

Various internal and external factors, including drugs, nutrients and metabolites, as well as non-molecular influences, such as radiation, etc., induce alterations in the stability of proteins in cell lysates, living cells and organisms. These alterations can be probed on the proteome-wide scale by applying a stability-modifying agent, such as elevated temperature (Savitski et al., 2014), proteolytic enzyme (Lomenik et al., 2009; Piazza et al., 2018), chaotropic agent, such as urea (Park and Maqusee, 2005), or salt (Vedadi et al., 2006). The agent is typically applied to a varying degree in a step-wise manner, and the fraction of the proteome remaining stable or, alternatively, the fraction becoming unstable, is analyzed. The protein stability can be assessed by, e.g., measuring the fraction of molecules remaining soluble at given conditions. The obtained information can be interpreted as drug binding to protein targets (Becher et al., 2016; Dart et al., 2018), as well as protein-protein docking, protein-small molecule interactions, or post-translational modifications (Huber et al., 2015; Becher et al., 2018; Dai et al., 2018; Saei et al., 2018).

One such popular method of monitoring the changes in protein stability is thermal proteome profiling (TPP) (Savitski et al., 2014), that has translated onto a proteome-wide scale the targeted approach of cellular thermal shift assay, CETSA (Molina et al., 2013), which, in turn, is based on a well known concept of protein melting temperature shift, widely used in drug discovery and development. Changes in protein’s physicochemical properties due to temperature variations have previously been applied to test interactions of this protein with other molecules using as a read-out fluorescence (Lo et al., 2004; Nielsen et al., 2007), calorimetry (Bruylants et al., 2005), differential scanning calorimetry (Brandts & Lin, 1990) or mass spectrometry (West et al., 2012). For instance, Garbett et al. (Garbett et al., 2009) have used differential scanning calorimetry of the unfractionated plasma, attributing changes in signature thermograms not to changes in the protein concentrations, but to interactions of these proteins with small molecules and peptides.

TPP is not restricted to the detection of protein-compound interactions (e.g., Park et al., 2017; Massey, 2018; Miettinen et al., 2018; Türkowsky et al., 2018), but has also been used for probing protein stability (Mateus et al., 2017; Volkening et al., 2018), its changes in cells (Becher et al., 2018; Dai et al., 2018) and bacteria (Mateus et al., 2018). Recently, we have developed based on TPP the new method of System-wide Identification of Enzyme Substrates by Thermal Assay (SIESTA) (Saei et al., 2018).

In a typical TPP experiment, aiming at profiling the proteome changes due to added small-molecule drug, both drug-treated and untreated (control) biological systems are incubated at N_t_≥10 different temperature points, after which the systems are lysed, and insoluble proteins (typically, the molecules that lost their native structure due to thermal unfolding) are removed by ultracentrifugation. The proteins remaining soluble are reduced, alkylated and then digested by trypsin, upon which an isotopic label (e.g., tandem mass tag, TMT) is chemically attached. The labeled digests corresponding to different temperature points are then multiplexed into a single mixture, which is analyzed subsequently by LC-MS/MS. A straightforward LC-MS/MS analysis of a TMT mixture provides identification and quantification of ≤5000 proteins. In order to increase the depth of the proteomics analysis to 5000-10,000 proteins, the mixture is typically separated into 8-24 fractions, with each fraction being analyzed by LC-MS/MS individually.

To thus obtained relative abundances of thousands of proteins, sigmoid curves are then fitted one by one, with the middle point of the curve corresponding to the melting temperature Tm of a given protein (Figure 1a). For each protein, the melting temperature shift ΔTm induced by the drug or other factor of interest is then determined as the difference between the Tm values with and without the acting factor. Since for obtaining statistical significance for a given ΔTm value the whole analysis has to be repeated at least twice (two replicates), a minimal TPP experiment with 8 fractions requires 2×2×8=32 LC-MS/MS runs, with each run lasting 1.5-2.0 h.

**Figure 1.**
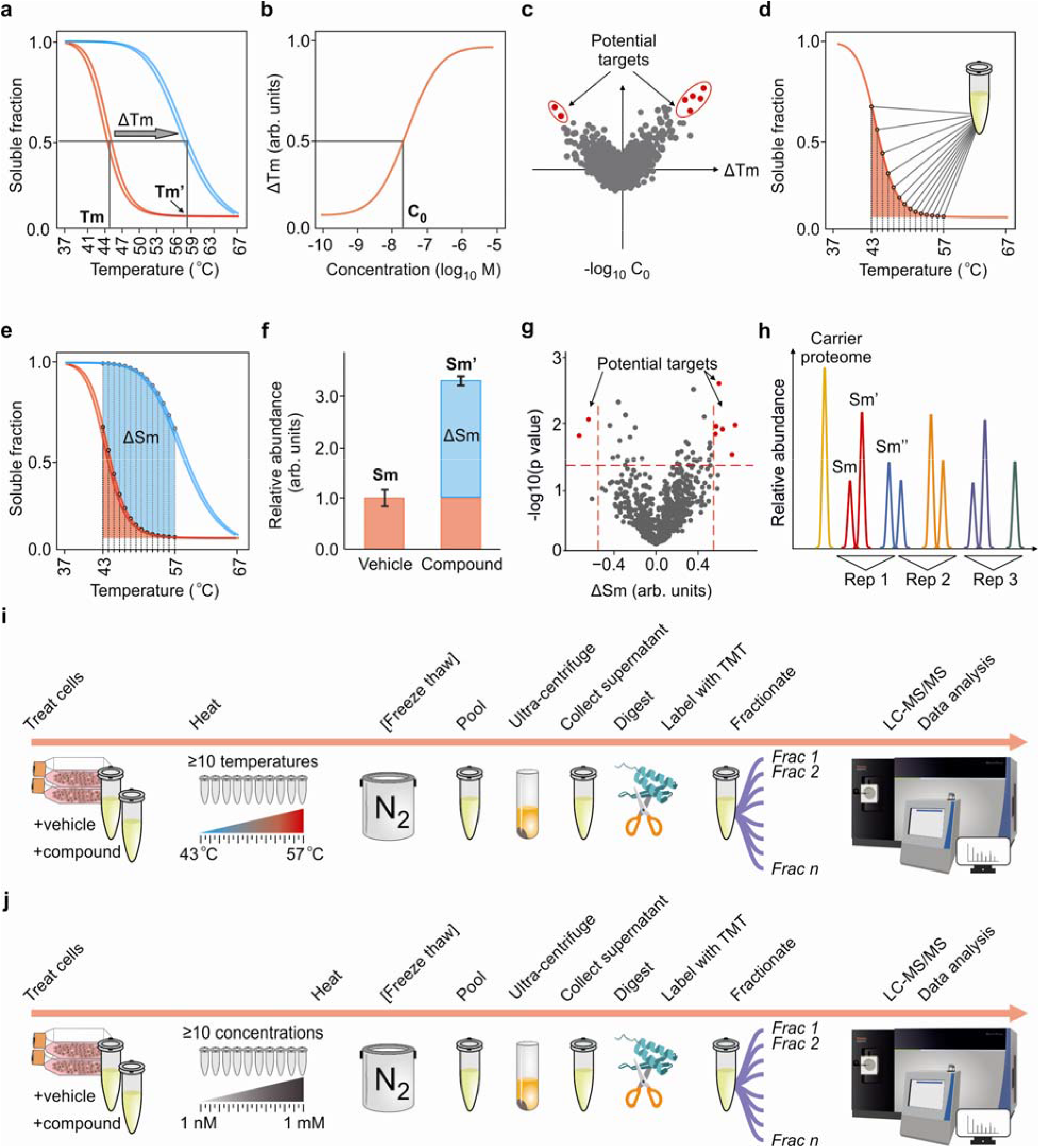
Schematics of the TPP and PISA assays. Thermal proteome profiling’s concept of sigmoidal curve fitting to **(a)** temperature scanning data for treated (blue) and untreated (red) samples, curve fitting and obtaining ΔTm, and **(b)** the same for concentration scanning data, with C_0_ determination. **(c)** Combination of the temperature and concentration domain data into a 2D plot and determination of the most probable protein candidates with large absolute ΔT shifts and low C_0_ values. **(d)** PISA concept of pooling together individual samples corresponding to different temperature points and thus hardware integration of the melting curve without detailed determination of its shape. **(e, f)** measuring ΔSm as a difference between the integral abundances of the protein in the treated and untreated samples. **(g)** Volcano plot of the 1D PISA results and determination of target candidates. **(h)** Example of a TMT-labeling scheme for a triplicate 2D PISA experiment. **(i)** 1D PISA experimental workflow, for each replicate sample: each sample in split in N_t_ ≥ 10 equal parts, each part being heated at a specific elevated temperature. If cells are used, they are then lysed by free-thaw cycles. Then the samples are pooled together before ultracentrifugation; collected supernatant is then digested and TMT-labeled, which is followed by fractionation and high resolution LC-MS/MS analysis with subsequent data processing. **(i)** The same for the second dimension, concentration-dependent experiment performed at a fixed elevated temperature.

To determine which proteins are most affected by the agent, the proteins are sorted by their ΔTm values, with positive ΔTm corresponding to stabilization, and negative - to destabilization by the factor. The p-value of a non-zero ΔTm can be determined by a two-tailed Student’s t-test using the two replicate measurements, or by an equivalent statistical method.

Often, such a 1D TPP analysis does not provide sufficient specificity to uniquely identify the protein most affected by the factor or even confine it to a short list of 3-5 most probable candidates. For specificity increase, a second dimension of analysis can be added, where the drug concentration (or factor intensity) is varied from zero to some maximum value (usually, 5-10 times higher than the IC_50_), while the temperature is fixed (Savitski et al., 2014). A sigmoid curve fitting can also be performed in the concentration domain, similar to the temperature domain (Figure 1b). The measured parameter in the second dimension is the concentration C_0_ (or pEC50, as in Savitski et al., 2014) at which the drug induces in a given protein thermal shift amounting to half of the ΔTm value. To obtain good curve fitting, a similar number of discrete concentration points N_c_ is needed as temperature points, preferably N_c_≥10. After determination of C_0_, proteins exhibiting largest absolute ΔTm values and the lowest C_0_ values, are selected as potential drug targets or mechanistic proteins responding to the factor (Figure 1c).

The above 2D TPP analysis is very powerful, but extremely resource consuming. A typical 2D procedure covering the whole TPP temperature range or at least the most useful half of it (usually, between 43 °C and 57 °C, encompassing the Tm values of most proteins) would require preparing and running of at least (10×32)/2=160 individual LC-MS/MS analyses. This may take more than two weeks of LC-MS/MS instrumental time, including the necessary blank runs between distinct groups of samples. Even with modern, reliable instrumentation, the risk of an unexpected stop or instrumental failure during such an experiment is non-negligible. The time and cost of such an analysis are the limiting factors in wider use of this powerful method in molecular biology and drug discovery.

The current 2D TPP approach possesses another limitation, which is the large number of cells needed for each samples (≈10^6^). Growing and handling ≥1.5•10^8^ cells per experiment is a challenge for any cell culture facility, since all cells need to be in a nearly identical biological state, as even small changes in the environment during cell growth can significantly affect the abundances of cellular proteins (Sabatier et al., 2018).

The curve fitting procedure to obtain Tm values presents another challenge. Some proteins increase (or appear to increase) their solubility with temperature, as do most molecules, and only upon reaching a significantly elevated temperature their structure starts to unfold, which finally results in solubility drop. The melting curves of such proteins can exhibit bumps, bimodal behavior or other unexpected features that can result in a low fitting score (Franken et al., 2015). Indeed, although the melting of proteins is usually considered to be a two-state transition from a defined folded native structure to a random coil, intermediate structures are often present (Biltonen & Freire, 1978; El-Baba et al., 2017). Moreover, some proteins, e.g. ribosomal units, are engaged in strong noncovalent complexes that fall apart only at significantly elevated temperatures. A sigmoidal curve fails to fit properly the melting behavior of many such proteins. Recently introduced non-parametric analysis of TPP data is more robust against deviations from the expected sigmoid shape (Childs et al., 2018), but it alleviates the problem of poor fitting only partially. As a result, problematic proteins are usually discarded from the final protein list, which increases the risk of false negative identifications (misses).

In order to solve or drastically reduce the impact of the above problems on the analysis results, we developed the Proteome Integral Stability Alteration (PISA) assay which achieves dramatic reduction in both analysis time and sample consumption by taking the following steps:

1. In a 1D PISA assay, two protein samples per replicate are analysed, one with the factor (drug) applied and another one – without the factor. For each of these two samples, the protein mixtures corresponding to N_t_≥10 individual temperature points are prepared, commonly 15 temperature points sampled at 1.0-1.5 °C starting from 43 °C. However, instead of labeling each of these samples by individual TMT following centrifugation and digestion, as in TPP, these protein mixtures are instead pooled together. The integral sample is then centrifuged, digested and labeled by a single TMT (Figure 1d). Thus a standard TMT-10 multiplexing scheme can combine 5 drug-treated and 5 untreated samples, which upon separation into 8 fractions will require less than a day of LC-MS/MS instrumental time to obtain the depth of >5000 proteins.
2. In data processing, instead of fitting to protein abundances at different temperature points a sigmoid curve and extracting Tm as in TPP, the read-out in 1D PISA is the protein abundance Sm in the pooled sample. This abundance represents the integral of the melting curve, independent of its actual shape (Figure 1e). If Sm is the read-out for the untreated sample and Sm’ is the corresponding value for the treated sample, then the PISA analogue of ΔTm is the function Ft(Sm, Sm’) that combines these two read-out values. In the simplest form, F_t_ = Sm’-Sm (Figure 1f), but we also found F_t_ = Sm’/Sm useful. The p-values can be determined by, e.g., Student’s t-test, and a volcano plot, as in expression proteomics, highlights the candidate proteins (Figure 1g).
3. In a 2D PISA assay, three samples are measured per replicate analysis (Figure 1h). The first two samples are the same as in 1D PISA (Figure 1i), i.e., one obtained without the drug (zero concentration), and another with the maximum drug concentration. These two samples provide Sm and Sm’ as read-outs, from which one obtains F_t_, as in 1D PISA. The third sample is a pool of the protein mixtures where intermediate drug concentrations are used (Figure 1j), and it provides the protein abundance Sm” as a read-out, Sm” being the integral of the concentration-dependence curve. The analogue of TPP’s read-out C_0_ is the function F_c_(Sm, Sm’, Sm”), which will be described below. Thus the 2D PISA assay provides two independent output parameters, F_t_ and F_c_, which can be plotted against each other in a 2D plot. Similar to 2D TPP, the proteins of interest usually combine extreme values of F_t_ and the maximum F_c_ values (Figure 1c). The standard TMT-10 labeling set can multiplex three 2D PISA replicates, with one TMT channel remaining vacant. This vacancy can be filled by an untreated proteome, which can be used for normalizing the S_m_ values (Figure 1h). According to our simulations, such a normalization can somewhat improve the precision. Alternatively, the vacancy can be filled by an untreated proteome obtained using detergent for enhanced protein extraction. Conventional TPP protocols tend to avoid detergents, as they affect protein solubility (Franken et al., 2015; Seashore-Ludlow & Lundbäck, 2016). This avoidance results in underrepresentation of less soluble proteins in TPP; inclusion of the detergent-assisted untreated proteome may cure this deficiency. The untreated proteome will play here the role similar to the carrier proteome in the single-cell proteomics approach introduced by Budnik et al. (Budnik et al., 2018).
4. As the simulation analysis presented below shows, there is a good linear correlation between the Sm values and the corresponding melting temperatures Tm, as well as between Sm’ and Tm’ values, provided the melting curves are sigmoidal. Thus, from the Sm and Sm’ data, one can extract via modeling the estimates of the Tm and Tm’ values. Moreover, there is a good linear correlation between ΔTm and F_t_ under the same sigmoidal curve assumption. Similarly, there is a good linear correlation between C_0_ and F_c_. Therefore, in PISA the information on these parameters in not lost, and can be derived through modeling if needed.

## Simulations

### Ab initio simulation of 2D PISA

As a theoretical proof of principle, we simulated in Excel the melting curves of 1000 proteins, with the melting temperatures Tm chosen randomly in the range from 42 °C to 57 °C. N_t_=16 temperature points between 37 °C and 67 °C with a 2 °C step were chosen. Sigmoidal melting curves were simulated by calculating the relative intensity I(T) for a given temperature T as:

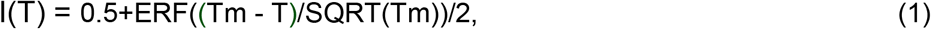

where ERF is the error function, and SQRT - the square root function. Examples of thus simulated melting curves are given in Figure 2a.

**Figure 2.**
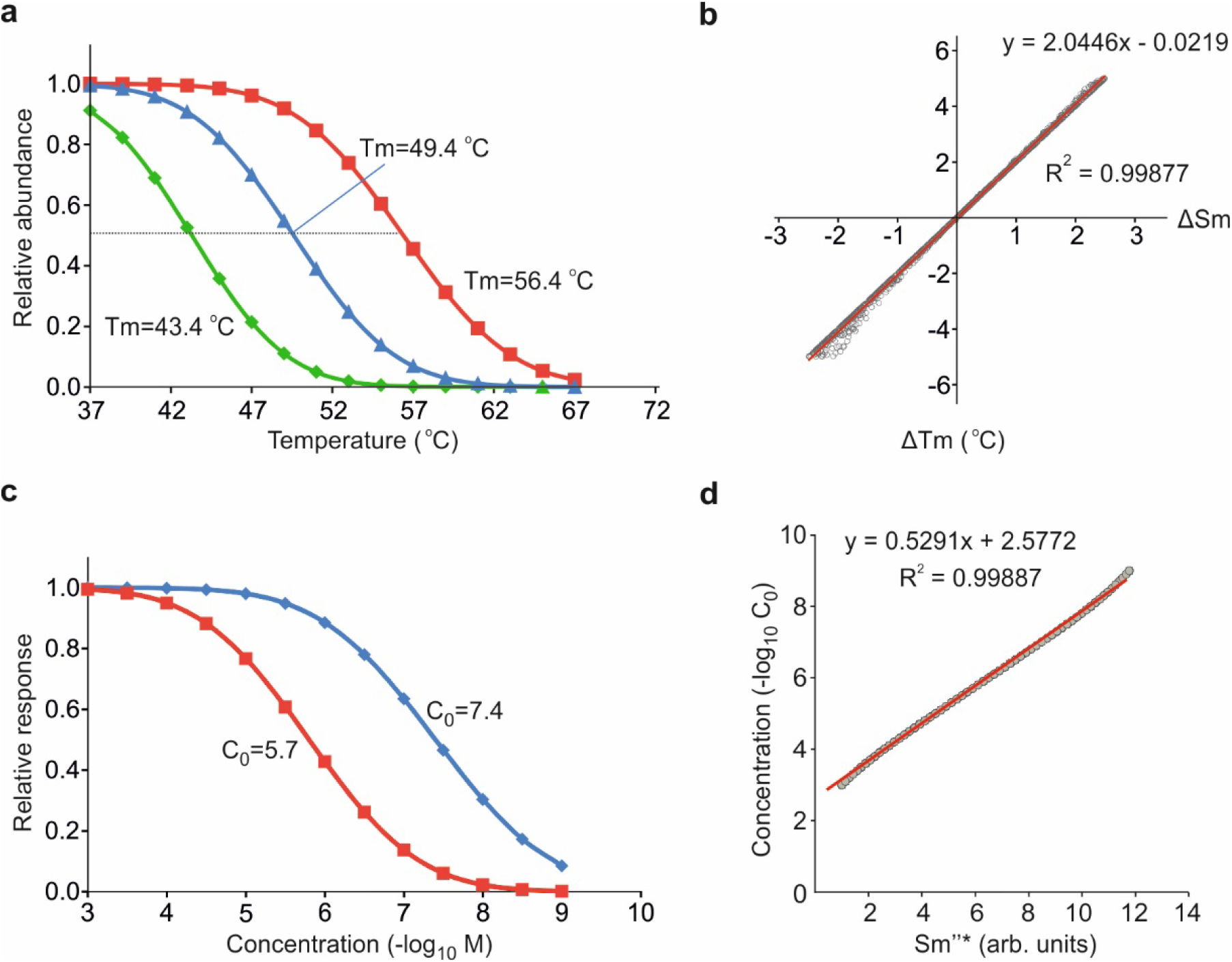
*Ab initio* simulation of 2D PISA. **(a)** Simulated melting curves. **(b)** Correlation between the simulated F_t_ = Sm′- Sm and ΔTm values. **(c)** Simulated response of the thermal protein stability shift to different drug concentrations. **(d)** Correlation between the calculated Sm”* values and C_0_ concentration.

The measured signal Sm was calculated as the sum of I(T) values for all temperature points. When the proteins were ranked by Tm and, separately, by Sm, the ranks were found to be exactly the same, *validating the hypothesis that Sm is a suitable proxy for Tm*.

The drug-induced melting temperature shifts ΔTm were simulated as random values in the range between −5 °C and +5 °C, and Tm’ values were calculated as Tm’ = Tm + ΔTm. The corresponding melting curve intensities I’(T) were calculated by (1) with Tm’ being used instead of Tm. The Sm’ values were obtained as the sums of all individual I’ values. F_t_ values were obtained as Sm’ – Sm, and correlated with the corresponding ΔTm values (Figure 2b). Excellent if not perfect (R^2^ > 0.998), this correlation proves that linear function is a suitable approximation. Knowing the parameters of the linear regression, one can derive the model Tm value from the measured Sm data.

The relative response R(C) of the thermal protein solubility shift to different drug concentrations C was modeled as a sigmoidal curve:

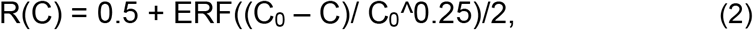

where C is a given concentration point, and ^ is the power function. Examples of the response function for two different C_0_ values are given in Figure 2c.

The measured read-out Sm” is simulated as Sm” = SUM(Sm + R(C)*(Sm’ - Sm)). The reduced parameter Sm”* is extracted from the measured values as

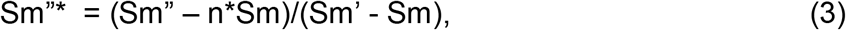

where n is the number of concentration points used. Again, the protein rank by Sm”* turned out to coincide with the rank by C_0_, confirming that the former parameter is a suitable proxy of the latter. An excellent linear correlation (R^2^ > 0.998) was found between Sm”* and −log_10_(C_0_) (Figure 2d). Thus, using linear regression, one can extract the model C_0_ values from S”* data.

These simple *ab initio* simulations provided theoretical proof of principle for the 2D PISA method. Not only the ΔTm and C_0_ protein ranks are preserved in the PISA output parameters, but there is a linear correlation between these parameters and the underlying ΔTm and C_0_ values.

### Simulation of 1D PISA output from experimental TPP results

Another proof of principle for the PISA approach was derived from the conventional TPP data, in which the abundances corresponding to different temperature points were added together to simulate the PISA read-out. From the published TPP datasets where both cells and cell extracts were treated with Dasatinib (Savitski et al., 2014), we calculated the Sm and Sm’ values as sums of protein abundances at all temperature points between 41 °C and 59 °C. The average correlation between the two replicates of the ΔTm values of the proteins surviving all stringent TPP filters for curve fitting were 0.903, while the average correlation between the corresponding ΔSm values was higher, 0.946. The better correlation in PISA corresponds to higher precision of the Sm calculation method compared to curve fitting in TPP. As a result, using the same statistical criteria for significance, there were 251 statistically significant proteins in TPP of cells and lysates, while there were 259 significant proteins in PISA (Figure 3a-b and d-e, respectively). At the same time, there was an excellent correlation between the replicate-averaged ΔTm and ΔSm values, R=0.947 for cells and R=0.987 for lysate (Figure 3c and f, respectively). These correlations were actually higher that those between the two ΔTm replicates in the original datasets, which were R=0.849 and R=0.931, for cells and lysate, respectively.

**Figure 3.**
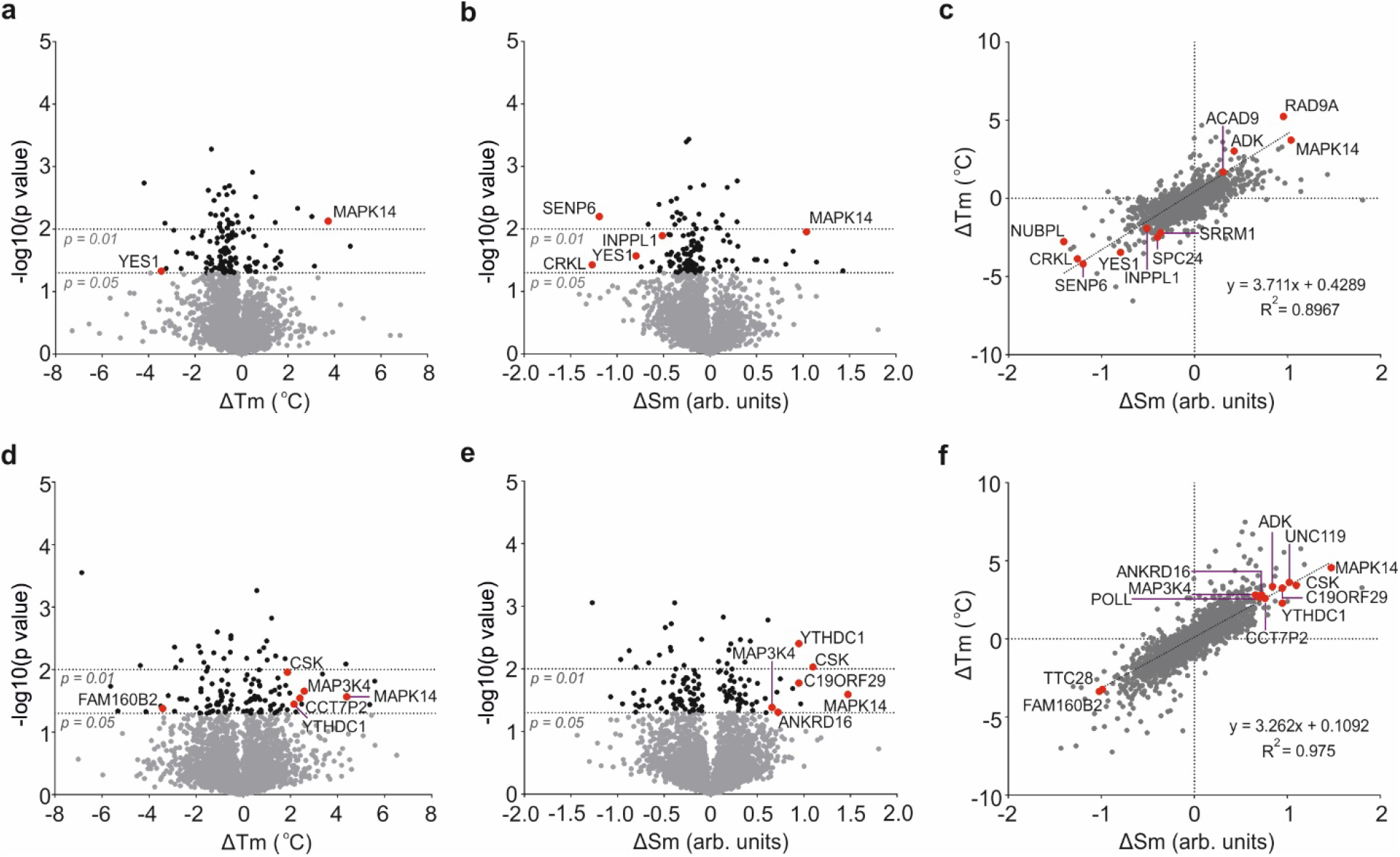
Simulation of PISA data analysis based on published TPP datasets. **(a)** Volcano plot (ΔTm, p) for TPP data of 5 μM Dasatinib treatment of cells. Proteins with significant (p<0.05) changes are shown in black. **(b)** The corresponding volcano plot (ΔSm, p) of the calculated PISA results. **(c)**. Correlation between the average ΔTm and average ΔSm values for Dasatinib treatment of cells. **(d-f)** Same as **(a-c)** for cell lysate extracts, respectively. Data are taken from (Savitski et al., 2014).

## Experimental

### 1D PISA assays on MTX and 5-FU

For purely experimental proof of principle, the PISA assay was performed on cells using two cell lines (lung carcinoma A549 and kidney carcinoma A498) and two drugs with well-known targets. In particular, we used the folate and nucleoside analogs methotrexate MTX and 5-fluorouracil (5-FU), known to inhibit, respectively, DHFR, an enzyme involved in the tetrahydrofolate synthesis, and TYMS, a key enzyme in de novo synthesis of thymidylate (Vincente et al., 2013; Qiu et al., 2017; Rajagopalan et al., 2002; Visentin et al. 2012; Wyatt et al. 2009). These drugs and their targets have already been subjects of CETSA investigations using antibody-based targeted protein detection, with the targets showing an increased stability after drug incubation (Jafari et al., 2014; Almqwist et al., 2016). The identification of DHFR and TYMS as targets of MTX and 5-FU, respectively, was also performed by Functional identification by expression proteomics (FITExP) (Chernobrovkin et al., 2014), a proteome-wide MS-based proteomics method for drug target deconvolution which is orthogonal to thermal shift approaches.

The first step in the PISA assay was to measure IC_50_ values of MTX in A549 cells and 5-FU in A498 cells, determined as drug concentrations causing 50% growth inhibition at 48 h. The obtained IC_50_ values were 1 μM for MTX in A549 and 17.5 μM for 5-FU in A498 cells. The subsequent steps were as in the workflow in Figure 1h. For treatment, we used a drug concentration corresponding to 5-10 times the IC_50_ value, and vehicle (solution without the drug) as a control. All experiments on both cells and lysates were performed in 5 biological replicates. The lysates were treated at N_t_=15 temperature points for 3 min each, and then pooled together after allowing precipitation at room temperature for 6 min and before centrifugation and digestion. Following TMT10 labeling of the digests, the samples were pooled and separated into 24 fractions by high pH reverse phase chromatography. The LC-MS/MS analysis was performed on an Orbitrap Q Exactive HF system. Protein identification and relative quantification was performed by as conventional in shotgun proteomics.

For MTX in A549 cells (Figure 4a), both DHFR and TYMS were clearly determined as by far the biggest positive outliers in the ΔSm distribution. In lysate, only DHFR was an outlier (Figure 4b), while ΔSm value for TYMS was very small. This result confirmed DHFR to be the primary target of MTX and TYMS to be a secondary target that binds a metabolized form of MTX (Qiu et al., 2017). Similar results were obtained for 5-FU that also binds to TYMS after metabolic modification (Longley et al., 2003): while in A498 cells TYMS is a clear positive outlier (Figure 4c), in a lysate the ΔSm value for TYMS was not significant (Figure 4d).

**Figure 4.**
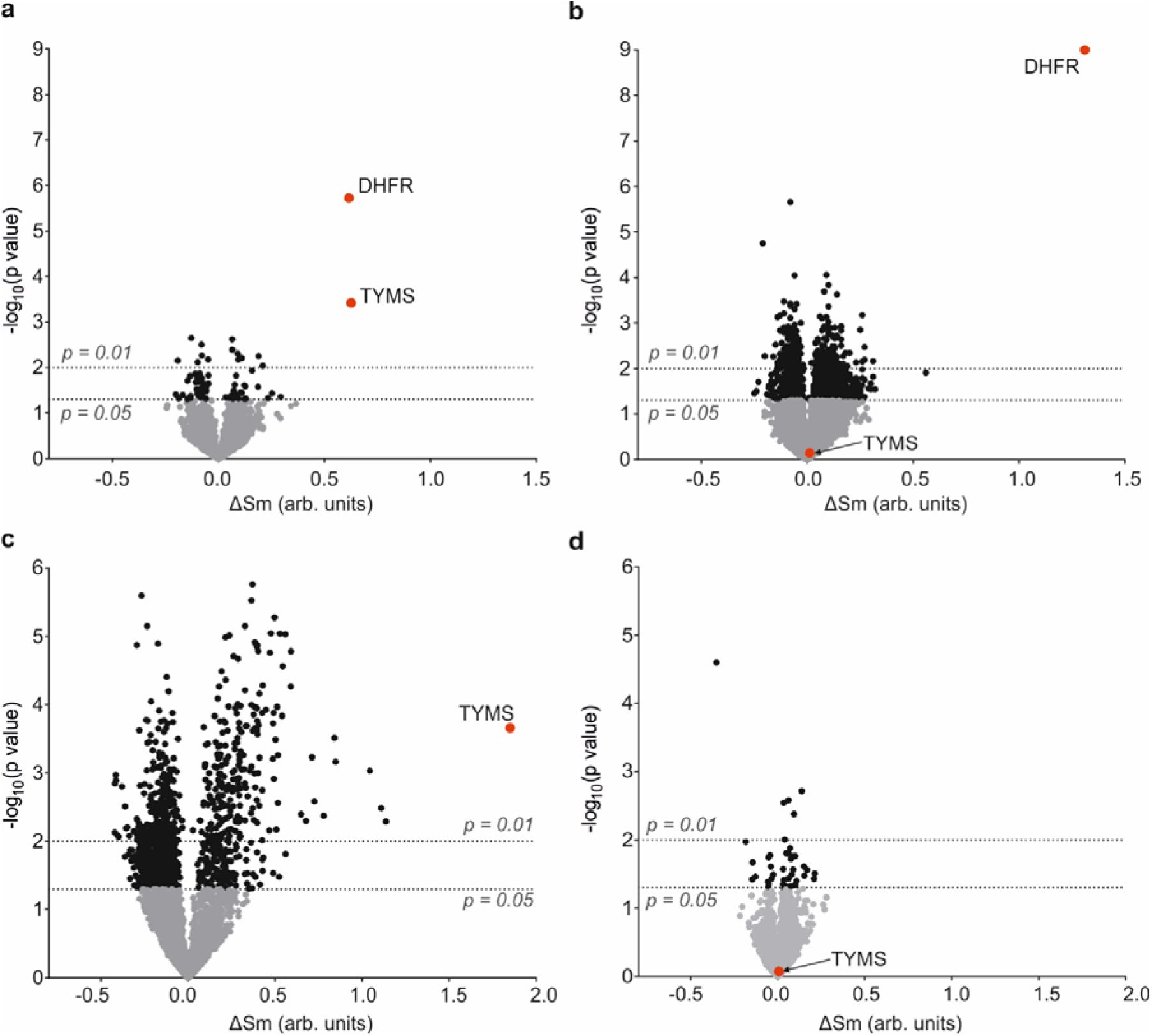
Volcano plots of experimental PISA assay results. Proteins with significant (p<0.05) changes are shown in black; known targets - in red. **(a)** A549 cells treated with MTX. **(b)** Same for cell lysate. **(c).** A498 cells treated with 5-FU. **(d)** Same for cell lysate.

### 1D PISA assay on 9 drugs

To demonstrate the unique analytical power of PISA arising from great reduction of the sample number, we treated A549 cell lysate with 9 drugs (Nutlin, Tomudex, Floxuridine, 8-azaguanine, Topotecan, Bortezomib, Dasatinib, Gefitinib, and Vincristine) at 10 μM for 45 min as well as vehicle. These 9 drugs have previously been used in deep-proteome FITExP analysis (Saei et al., 2018). Then we performed PISA analysis, multiplexing the 9 drugs and control into a TMT-10 sample in a biological triplicate, separating each replicate into 24 fractions and analyzing them by LC-MS/MS. The 72 analyses took less than a week of instrumental time, while with TPP the same effort would require two and a half month, a prohibitively expensive enterprise for most research groups. The analysis yielded 7200 proteins quantified in all replicates, and allowed us to apply the specificity concept previously used only in FITExP and SIESTA – namely, contrasting, for every protein, the ΔSm value for any particular drug to those for all other drugs. This was done by the OPLS-DA method. As an example, Dasatinib that targets kinases showed many kinases specifically stabilized or destabilized. To increase the analysis specificity, Gefitinib that has similar targets was removed from the dataset (Figure 5a).

For the first time, it became possible to compare the data on specific expression (FITExP) with those on specific thermal shift (PISA) on 5600 common proteins and prove the orthogonality of these two methods. As an example, the Floxuridine target TYMS shows elevated expression as well as positive stabilisation; it is an outlier in both types of analysis when Tomudex that has the same target is removed (Figure 5b).

**Figure 5.**
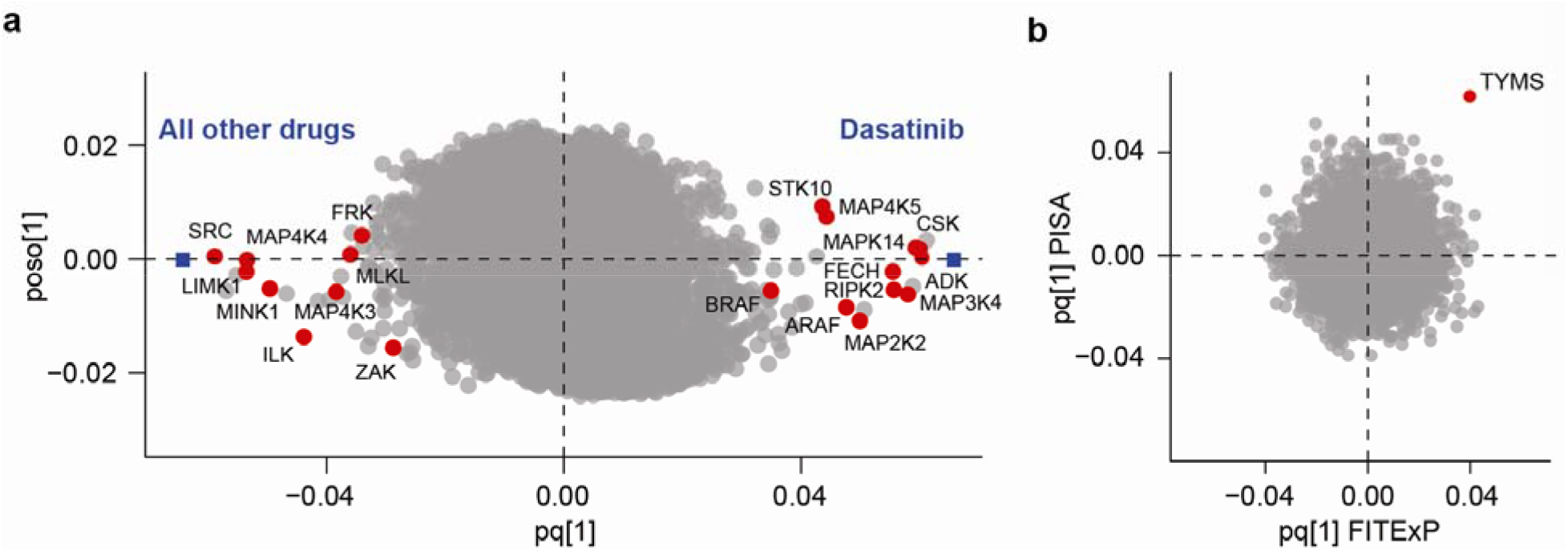
Results of a single PISA assay for 9 drugs. **(a)** OPLS-DA plot of Dasatinib data contrasted with all treatments, except Gefitinib. Known targets (kinases) are shown in red. **(b)** Comparison of FITExP results with PISA assay data on Floxuridine treatment for 5600 common proteins. The Floxuridine target TYMS shows most elevated expression as well as highest positive stabilisation.

## Discussion

### Specificity increase

The specificity increase in PISA compared to TPP can come from three sources. First, it is the possible use of a larger *number* of temperature points N_t_, as this incurs no additional cost in terms of LC-MS/MS instrumental time. Larger N_t_ with the same temperature *range* means smaller errors associated with the discrete character of measurements, and more accurate capturing of the behavior of proteins with steep melting curves. The use of more temperature points will also mitigate the error arising due to statistical noise, e.g., single point outliers. The only limitation on N_t_ is practical. We tested N_t_=20 temperature points and received lower p-values for known target proteins than with N_t_=10. Note that, for increased dynamic range of the readout, the temperature range can be narrowed to the region of the most significant solubility changes, excluding both the lowest and the highest temperatures where the difference between the treated and untreated samples is small.

The second reason for the specificity increase in PISA compared to TPP is the use as a readout of the statistically robust integral under the melting curve Sm instead of the curve-fitting parameters that are subjects to various statistical uncertainties. Also, since all samples corresponding to various temperature points are pooled together before lysis (when intact cells are drug-treated) or ultracentrifugation (when lysates are treated), all the downstream procedures, such as reduction/alkylation, digestion and TMT labeling, are performed on a single pooled sample, which reduces the experimental errors and thus improves the p-values. Of course, when different drug concentrations are used, samples have to be pooled after centrifugation and supernatant collection.

The third reason is the larger number of replicates that can be analyzed in practice. We mentioned above that a standard TMT-10 labeling scheme allows for simultaneous analysis of five replicates of both treated and untreated samples, while in published TPP studies we found no more than three replicates. The much larger statistical power of the five-replicate experiment allows one to identify with high significance even proteins with tiny thermal shift.

### Throughput increase

With the same number of replicates, 1D PISA analysis provides a throughput increase by a factor of N_t_ compared to TPP, while 2D PISA gives an increase of N_t_·N_c_. With both Nt and Nc being of the order of 10, the throughput increase in 2D PISA can reach two orders of magnitude.

### Sensitivity increase

One of the limiting factors in TPP is the minimum volume (usually, ≈10 μL) of lysate allowing for reliable supernatant collection after ultracentrifugation of the thermally treated sample. In 1D PISA, where the samples are merged before centrifugation, reduction of the minimal volume per sample is by a factor of Nt. Similarly, in 2D PISA, where the samples for different drug concentrations are merged before thermal treatment, the overall reduction of sample volume is N_t_·N_c_ times.

### Cost reduction

The costs of a PISA experiment arises from the use of biological materials, chemicals (e.g., TMT10 labeling reagents), labor for sample treatment and preparation for the LC-MS/MS analysis, as well as the LC-MS/MS analysis itself. The reduction in the volume of biological material (mainly, cells) is similar to the above increase in sensitivity. In drug discovery, the cost of an experimental drug can be quite substantial, and thus the cost reduction in PISA can be high. Besides, in our recent TPP-based method of SIESTA (Saei et al., 2018), where a recombinant enzyme is added to a cell lysate together with a co-factor, the cost of a recombinantly produced and purified enzyme with validated activity can exceed the cost of the LC-MS/MS part of SIESTA, scaling up with the required enzyme amount. Therefore, SIESTA is one of the analysis types that will greatly benefit from the use of PISA instead of TPP. Additional cost reduction comes from the TMT reagents and LC-MS/MS instrumental time, which are usually the most expensive items in TPP. Paradoxically, labor and chemicals become the dominant cost items in PISA while the LC-MS/MS instrumental time becomes a lesser component.

### Applicability area

The main idea behind PISA – the use of the integral under the curve instead of the curve shape parameters – can be applied in many analytical methods were curve fitting is employed to probe protein stability or solubility, e.g. in limited proteolysis combined with MS (Leuenberger et al., 2017), in the use of urea or other chaotropic agents, pressure or high (low) pH values, or high (low) salt concentrations.

